# Response of Serum-isolated Extracellular Vesicles to Focused Ultrasound Blood-Brain Barrier Opening

**DOI:** 10.1101/2024.12.17.629012

**Authors:** Alina R. Kline-Schoder, Fotios N. Tsitsos, Alec J. Batts, Melody R. DiBenedetto, Keyu Liu, Sua Bae, Elisa E. Konofagou

**Author notes:** Corresponding authors: Fotios N. Tsitsos, Elisa E. Konofagou.

## Abstract

The blood-brain barrier (BBB) limits drug delivery to the brain and the movement of neurological biomarkers between the brain and blood. Focused ultrasound-mediated blood-brain barrier opening (FUS-BBBO) noninvasively opens the BBB, allowing increased molecular transport to and from the brain parenchyma. Despite being initially developed as a drug delivery method, FUS-BBBO has shown promise both as a neuroimmunotherapeutic modality, and as a way of improving neurological disease diagnosis via amplification of disease biomarker circulation.

Recently, the role of extracellular vesicles (EVs) in modulating the neuroimmune system and in improving biomarker detection has sparked research interest. However, despite their potential role in modulating FUS-BBBO-induced neuroimmunotherapy and their ability to improve biomarker specificity after treatment, the EV response to FUS-BBBO had not been extensively characterized prior to this study.

In this study, we investigated the effect of FUS-BBBO on EV concentration and content in the serum of mice and Alzheimer’s Disease (AD) patients. We observed a 164% increase in murine EV concentration one hour after treatment, as well as an increase in EV RNA associated with FUS-BBBO neuroimmunotherapy. Patient EV concentration also increased one hour after treatment and was dependent on the volume of BBB opening three days post-treatment. Furthermore, EV isolation was found to significantly enhance the amplification of AD biomarker detection by FUS-BBBO.

Overall, we present the first evidence of altered murine and AD patient EV concentration and content in response to FUS-BBBO, providing evidence of EVs’ role within FUS-BBBO neuroimmunotherapy as well as their utility in improving FUS-BBBO biomarker amplification.

## Introduction

The blood-brain barrier (BBB) is a virtually impermeable barrier between the blood and the brain that keeps the brain at homeostasis for neuronal firing. In addition to limiting the infiltration of neurotoxins and pathogens, the BBB limits both the delivery of drugs to the brain and the circulation of neurological disease biomarkers in the blood [1]. Focused ultrasound-mediated blood-brain barrier opening (FUS-BBBO) combines focused ultrasound with intravenously administered microbubbles to transiently and noninvasively open the BBB, tackling these two challenges [2], [3], [4].

Although initially designed to facilitate drug delivery through the BBB, FUS-BBBO has also been established as a neuroimmunotherapeutic treatment for neurological diseases and a method of amplifying the detection of neurological biomarkers [4], [5], [6], [7], [8], [9], [10]. As a neuroimmunotherapeutic, FUS-BBBO has been shown to reduce disease pathology and ameliorate disease-associated cognitive deficits in many neurological models ranging from Alzheimer’s Disease (AD) to depression [6], [11]. These effects coincide with brain macrophage modulation, increased neurogenesis, and increased synaptic plasticity [5], [12], [13]. FUS-BBBO amplification of neurological biomarkers has been primarily investigated in models of brain cancer, where reports have found increases in circulating cell-free DNA (cfDNA), as well as central nervous system (CNS) proteins, including Glial Fibrillary Acidic Protein (GFAP), in response to treatment [4], [7], [8], [9], [10].

Extracellular vesicles (EVs) are lipid vesicles responsible for cell transport and exchange. EVs have highly variable cargo, including proteins, carbohydrates, and/or coding and non-coding RNA (ncRNA). Due to their small size, there is a particular emphasis on ncRNA within EVs such as micro RNA (miRNA) and piwi-interacting RNA (piRNA). Both miRNA and piRNA regulate the mRNA transcription of target protein-encoding genes. There are publicly available databases of miRNA and piRNA target genes, which allow functional annotation of the up- and downregulated ncRNA targets [14].

EVs are reported to modulate the neuroimmune system, including maintenance and repair of the BBB, neurogenesis, and synaptic plasticity [15], [16], [17], [18], [19]. Due to their role in intercellular communication, EV isolation and characterization has emerged as a method of improving the specificity of biomarker detection. Recent metanalyses of biomarkers in AD have found that isolating EVs prior to quantifying protein load provides a more specific diagnosis [20].

There is preliminary evidence of *in vivo* FUS-BBBO increased neuronal EV concentration and *in vitro* FUS-BBBO increased neuroprotective EV concentration [21], [22]. Additional work in the periphery has identified a FUS-induced increase in anti-inflammatory EVs after treatment of arthritis [23], [24]. Given this preliminary evidence of FUS-BBBO affecting EV concentration and content, as well as their dual role in modulating the neuroimmune system, and as an emerging biomarker, we aimed to identify the effect of FUS-BBBO on EV concentration and content.

In this study, we explore the response of EVs in serum following FUS-BBBO in a mouse model and in a clinical trial with AD patients. In the mouse study, the concentration of EVs in serum, as well as their genomic and proteomic content are analyzed. To assess the role of EVs in modulating the neuroimmune response following FUS-BBBO, we investigated the effect of GW4869, a neutral sphingomyelinase inhibitor that blocks EV generation [25], on the restoration timeline of the BBB. Finally, to elucidate the potential of EVs in amplifying neurological marker detection, we collected blood samples from AD patients participating in our group’s clinical study on neuronavigation-guided FUS-BBBO and analyzed the protein content of EVs for important AD biomarkers. This study provides the first translational analysis of EV concentration and content after FUS-BBBO, providing evidence of both the EV role within the neuroimmunotherapeutic response and the potential use of isolating EVs to improve FUS-BBBO neurological biomarker accentuation.

## Materials and Methods

### FUS-BBBO in mice

All animal experiments were reviewed and approved by the Institutional Animal Care and Use Committee at Columbia University. Mice in the FUS-BBBO and microbubble (MB) sham groups were anesthetized with a mixture of oxygen and 1-2 % isoflurane (SurgiVet, Smiths Medical PM, Inc., WI), placed on a stereotaxic apparatus (David Kopf Instruments, Tujunga, CA) and their heads were immobilized and depilated to reduce acoustic impedance mismatch. Degassed ultrasound gel was applied on the head and a bath with degassed, deionized water was lowered on top of the head. The lambdoid suture was identified, and the transducer was positioned over it as previously described [26].

In mice that were treated with FUS-BBBO, a single-element, spherical-segment concave FUS transducer (center frequency: 1.5 MHz, focal depth: 60mm, radius: 30mm; Imasonic, France) that was driven by a function generator (Agilent Keysight 33220A, Palo Alto, CA, USA) through a 50-dB power amplifier (325LA, Electronic Navigation Industries, Rochester, NY, USA) was used to treat the two hippocampi. The center of the transducer held a pulse-echo ultrasound transducer (V320, center frequency: 7.5 MHz, focal depth: 52 mm, diameter 13 mm; Olympus NDT, Waltham, MA) that was used for alignment and passive monitoring of microbubble cavitation. The pulse-echo ultrasound transducer was driven by a pulser-receiver (5077 PR, Olympus, Waltham, MA, USA) which was in turn connected to a digitizer (Gage Applied Technologies, Inc., Lachine, QC, Canada). The transducer setup was attached to a three-dimensional positioning system (Velmex Inc., Lachine, QC, Canada). Each hippocampus was sonicated first for 10 seconds to obtain baseline cavitation dose and then again for 2 minutes for the experimental sonication. For all experiments, in-house synthesized, lipid-shelled microbubbles (average concentration: 8×10e8/mL, mean diameter: 1.4 μm) were manufactured according to previously published protocols [27], [28]. A bolus of 3 μL of microbubbles was diluted in 100 μL of sterile saline and was injected intravenously between the baseline and experimental sonications. The transducer was not triggered to treat MB sham mice; otherwise, the treatment was identical to that of the FUS-BBBO mice. Mice in the naïve group were not subjected to anesthesia, FUS or microbubble injections.

### Mouse Serum Collection

Mouse blood was collected from the submandibular vein without anesthesia the day before, and 1 hour after FUS-BBBO (Fig. **1A**). Mice were held by grasping the skin behind the head firmly. A 16-gauge needle was inserted into the submandibular vein and then removed. Less than 100 μL of blood was collected in heparin-coated serum-separating tubes (Ram Sciences). After collection, gentle pressure was applied to the site of the puncture in order to stop bleeding. After collection, blood was left at room temperature for 15-30 minutes to clot and then spun down at 3000 x g for 15 minutes. Serum was aliquoted to a separate tube and stored at -80 ^o^C for future processing.

**Figure 1.**
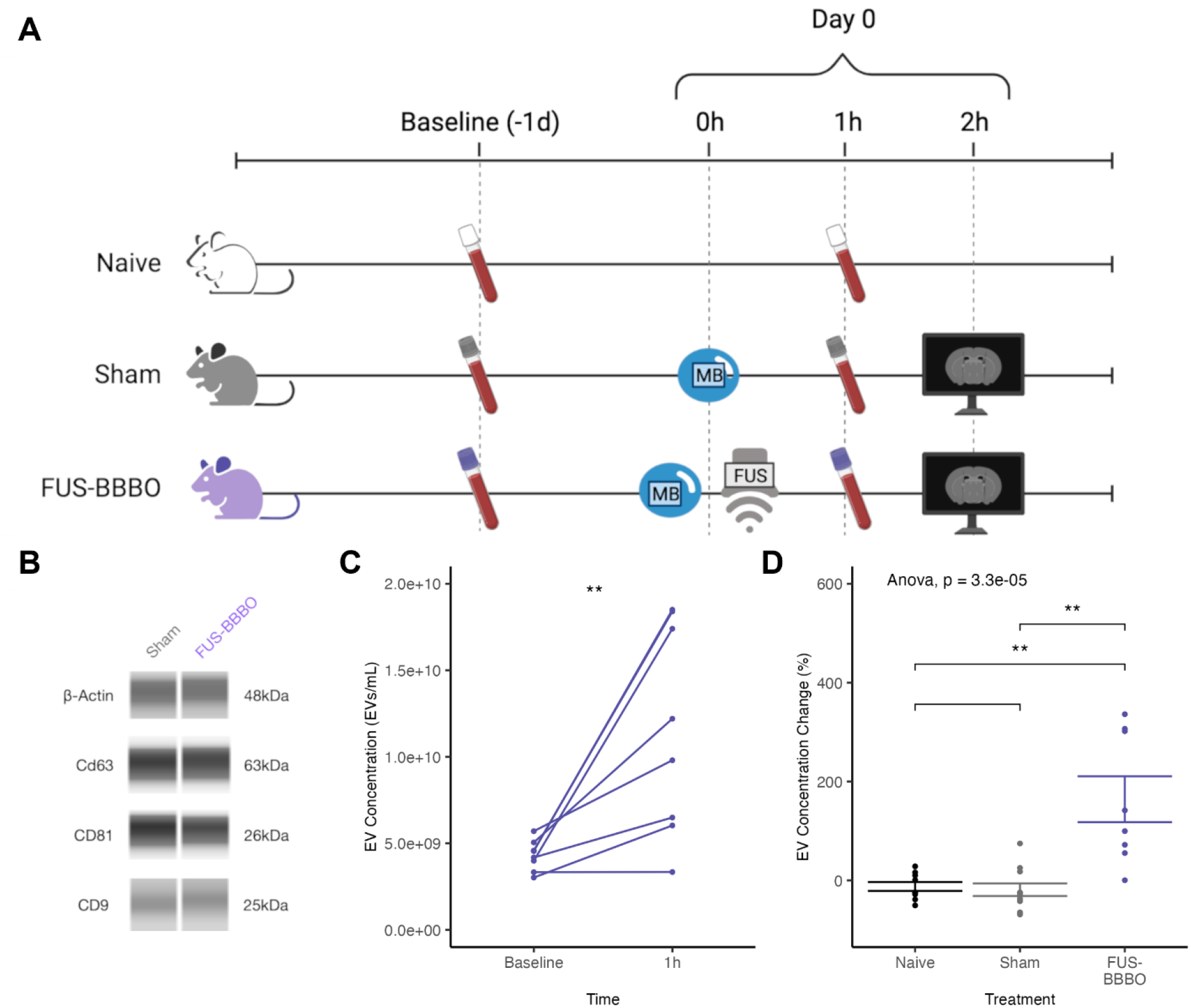
Extracellular vesicle concentration increases after FUS-BBBO in the serum of mice. **A)** Murine experimental timeline. Blood draws were taken for EV isolation at two time points: prior to treatment (baseline) and 1 hour after treatment (1 hour). Animals were randomly split into three treatment groups: FUS-BBBO, sham, and naïve. Mice in the FUS-BBBO group had 2 minutes of FUS applied bilaterally following MB injection at time 0. Animals in the sham group were injected with MB, but no FUS was applied. Mice from the FUS-BBBO and sham groups were injected with gadolinium (Gd), and a T1-Weighted MRI was performed less than 2 hours after the FUS-BBBO and sham procedures. Animals in the naïve group were anesthesia-, MB-, and FUS-naïve. **B)** Western blots of β-actin and EV tetraspanin markers CD9, CD63, and CD81 on representative sham and FUS-BBBO EVs from 1 hour post-treatment show successful EV isolation. **C)** EV concentration at baseline and 1 hour post-treatment in mouse serum. Paired t-test was performed and displayed on graph. **D)** Percent change in EV concentration at the 1-hour time point for all 3 groups; the FUS-BBBO group shows a significant change in EV concentration after 1 hour compared to the naïve and sham groups. One-way ANOVA followed by Bonferroni-corrected t-test between groups was performed. All statistics are displayed on the graph.

### Magnetic Resonance Imaging for Mice

Following treatment and blood draws, all FUS-BBBO and MB Sham animals underwent scanning with a 9.4 T MRI system (Bruker Medical, Boston, MA). Mice were intraperitoneally injected with 0.2 mL of gadodiamide solution (Omniscan^TM^, GE Healthcare, Princeton, NJ) exactly 30 minutes prior to scanning. Images were acquired using a T1-weighted 2D FLASH sequence (TR/TE 230/3.3 ms, flip angle: 70°, number of excitations: 6, field of view: 25.6 mm × 25.6 mm).

### GW4869 Drug Administration

For the EV inhibition study the drug GW4869 (MedChem Express, Monmouth Junction, NJ, USA), a neutral sphingomyelinase inhibitor, was used to block EV generation. Mice were separated in four groups: naïve, naïve+GW4869, FUS-BBBO, and FUS-BBBO+GW4869. A 2.5 mg/mL stock solution of GW4869 in dimethyl sulfoxide (DMSO) was prepared and stored at -20^°^C until the day of injection. Immediately before administration, the drug was further diluted in sterile saline to a final concentration of 0.25 mg/mL. The drug was administered intraperitoneally at a dose of 2.5 μg/g body weight per mouse.

To study the effect of EVs in the restoration of BBB following FUS-BBBO, mice in the FUS-BBBO and FUS-BBBO+GW4869 were monitored for five days after the sonication procedure. The FUS-BBBO+GW4869 group received injections of GW4869 before the FUS-BBBO treatment, and then on day 2 and day 4 after the procedure. The progress of BBBO restoration was monitored for both groups by contrast-enhanced T1-weighted MRI 2 hours after FUS-BBBO, and then on days 1, 3, and 5 post FUS-BBBO. Blood was collected on the day prior to FUS-BBBO, and then 1 hour and 1 day after FUS-BBBO, and always before the injection of gadolinium solution for MRI contrast.

To elucidate the role of EVs in the inflammatory response following FUS-BBBO, mice from all four groups were allowed to survive for one day after the FUS-BBBO groups were sonicated. Mice in the naïve+GW4869 and FUS-BBBO+GW4869 groups received one injection of the drug immediately before the FUS-BBBO procedure took place. Furthermore, BBBO was confirmed for the mice that underwent FUS-BBBO by contrast-enhanced MRI 2 hours after the sonication procedure. Mice from all groups had blood taken on the day prior to FUS-BBBO, 1 hour, and 1 day after FUS-BBBO. After the final blood draw, the mice were sacrificed by transcardial perfusion with cold 1X PBS and their hippocampi were separated for bulk RNA sequencing.

### FUS Treatment in AD Patients

Six Alzheimer’s Disease (AD) patients underwent neuronavigation-guided FUS-BBBO as part of our phase I clinical trial (NCT04118764). All methods used were approved by Columbia University’s Institutional Review Board and all participants provided informed consent prior to the FUS-BBBO procedure. The right frontal lobe was targeted and numerical simulations using the k-wave package in MATLAB were carried out prior to the experiment to estimate the power attenuation of FUS through the skull. A detailed account of the methods of FUS-BBBO in AD patients is given in our group’s clinical paper [29]. Briefly, a single-element, spherical-segment FUS transducer (H-231, Sonic Concepts, Bothell, WA, USA) was driven at a frequency of 0.25 MHz by a function generator (Agilent, Palo Alto, CA, USA) and a 50-dB power amplifier (Electronic Navigation Industries, Rochester, NY, USA) to emit FUS with pulse length 10 ms and pulse repetition frequency 2 Hz. A derated peak-negative FUS pressure of 200 kPa (mechanical index (MI) = 0.4) was applied for 2 minutes. A bolus of microbubbles (0.1 mL/kg, Definity, Lantheus) was intravenously injected at the start of the sonication. During the sonication, the cavitation dose was monitored by using a single-element transducer for the first four patients and an imaging array transducer (P4-2, ATL Philips) for the remaining two patients.

### Patient BBBO Volume Quantification

BBB opening volume was quantified from the contrast-enhanced T1-weighted MRIs, which were taken 2 and 72 hours after the FUS treatment using a 3 T system (Signa Premiere, General Electric, Boston, MA, USA) for the confirmation of the opening and closing of BBB respectively. For T1-weighted contrast enhancement, an intravenous injection of 0.2 mL/kg gadoterate meglumine (Dotarem®, Guerbet). The contrast-enhanced volume was quantified by subtracting the 72-hour contrast-enhanced T1 MRI from the 2-hour MRI and thresholding the subtracted image. The threshold was chosen automatically so that the average intensity within the opening volume was considerably higher than that of the surrounding area, with a 98% level of confidence assuming the Gaussian distribution of the intensity of the subtracted image.

### Extracellular Vesicle Isolation

The commercial ExoQuick (Systems Biosciences, Palo Alto, CA, USA) was used according to manufacturer instructions to isolate extracellular vesicles from both mouse and human serum. Briefly, 25 μL of mouse serum were diluted with 75 μL of 1X PBS before incubation with 22.5 μL of ExoQuick for 30 minutes at room temperature. For human samples, 200 μL of undiluted serum were incubated with 45 μL of ExoQuick for 30 minutes at room temperature. The samples were then spun down at 1500 x g for 35 minutes at 4^°^C, and the pellet (isolated extracellular vesicles) was resuspended in 100 μL or 200 μL of 1X PBS for the mouse and human samples respectively. The isolated EV suspensions were stored at -80 ^o^C until further processing.

### Nanoparticle Tracking Analysis

Extracellular vesicle concentration analysis was quantified by nanoparticle tracking analysis (NTA) on a NanoSight (NS300, Malvern Panalytical, Malvern, UK). 5 μL of isolated mouse EVs or 1 μL of isolated human EVs were further diluted into 1 mL of 1X PBS. This solution was then run through the Nanosight at a rate of 1000 μL/min, and the resulting image was captured and analyzed for particle concentration and size distribution.

### Multiplex Protein Quantification

For the samples from the clinical study, a Luminex multiplex assay was used to quantify proteins in the serum and isolated clinical extracellular vesicles (Luminex Corporation, Austin, TX, USA). Single procartaplex kits were purchased and combined to make a custom multiplex panel for analysis (Invitrogen, Waltham, MA, USA). Data were fit with a separate five-parameter logistic dose-response curve for each protein, and all curves had R^2^ greater than or equal to 0.95.

### Mass Spectrometry Protein Analysis

For the samples from the mouse studies, mass spectrometry proteomics was performed by Systems Biosciences on already isolated EVs. Briefly, Systems Biosciences lysed the isolated EVs in a gel-loading buffer, followed by gel-based extraction and trypsinization for peptide library creation for LC/MS ESI-TOF. Peptide signatures were then mapped to a database of known protein sequences. Peptide quantification across all four runs was loaded into R, normalized, and processed for differential protein expression utilizing the UniprotR package. All functional annotation was performed with the TopGO package.

### RNA Sequencing

RNA Sequencing was performed by Systems Biosciences on already isolated EVs and dissected frozen hippocampus tissue. RNA was isolated and quantified using Agilent Bioanalyzer Small RNA Assay before 75bp single-end read Next Gen Sequencing libraries were prepared with Qiagen small RNA library preparation and gel purification. Sequencing was performed on Illumina NextSeq with SE75 at an approximate depth of 10-15 million reads per sample. Reads were processed and aligned to the GRCm38 genome with Ensemble transcriptome annotation (GRCm38.p6) using CellRanger with default parameters. Count tables were loaded into R and underwent normalization and differential gene expression analysis with the edgeR package. piRNA and miRNA targets were extracted from piRNAdb and miRBase, respectively. All functional annotation was performed with the TopGO package.

### Western Blotting

RayBiotech performed western blotting using an automated capillary immunoassay method. Samples and reagents were loaded onto an assay plate and put into the western blotting machine. The sample was automatically loaded and separated by size while it traveled through the stacking and separation matrix. Then, the separated proteins are fixed with proprietary capture chemistry. Target proteins are identified with primary and secondary HRP-conjugated antibodies.

## Results

### Murine extracellular vesicle concentration increases after FUS-BBBO

Wild-type mice were separated into three groups – naïve, sham, and FUS-BBBO (Fig. **1A**). Animals treated with FUS-BBBO were intravenously injected with microbubbles (MB) and treated with 2 minutes of focused ultrasound bilaterally on the hippocampi, consistent with literature [26], [30]. Animals in the sham group were intravenously injected with MB but were not treated with focused ultrasound. Animals in all groups had blood drawn twice immediately before treatment (baseline) and 1 hour after treatment (1 hour). Animals in the sham and FUS-BBBO groups underwent contrast-enhanced T1-weighted MRI after the second blood draw to confirm the opening within the FUS-BBBO group and the lack of opening within the sham group. The methods detail the EV isolation and concentration quantification.

Western blots of the isolated EVs from representative sham and FUS-BBBO samples confirmed the successful isolation of the EVs via expression of marker tetraspanin proteins CD9, CD81, and CD63, as well as β-actin (Fig. **1B**). Nanoparticle tracking analysis (NTA) reveals that the EV concentration is significantly increased 1 hour after treatment compared to the baseline (Fig. **1C**). Furthermore, the average increase of 164% after FUS-BBBO is significantly higher than the percentage change in naïve and sham samples which averages near 0% (Fig. **1D**).

### FUS-BBBO alters murine extracellular vesicle protein and RNA load

Given the significant increase in EV concentration 1 hour after treatment, we performed whole genome RNA-sequencing and mass spectrometry protein identification (Systems Biosciences, Palo Alto, CA, USA) on isolated EVs from baseline and 1 hour. From each time point, two samples pooled from 3 animals underwent both processes. Differential gene expression analysis between 1 hour and baseline reveals significantly up- and downregulated protein-coding and non-protein-coding (ncRNA) RNA (Fig. **2A-C**). Upregulated protein-coding genes include proliferation-associated genes such as *Sox3* and inflammation-associated genes such as *Mapk12*. Downregulated protein-coding genes include tight-junction genes such as *Cldn11*. Differential expression of the protein identification finds many fewer significantly up and downregulated proteins (only 10 proteins compared to 900 genes). The most significantly upregulated proteins include immediate inflammatory response proteins such as *Lbp* and hemoglobin-associated proteins such as *Hbb-bs* (Fig. **2D-E**).

**Figure 2.**
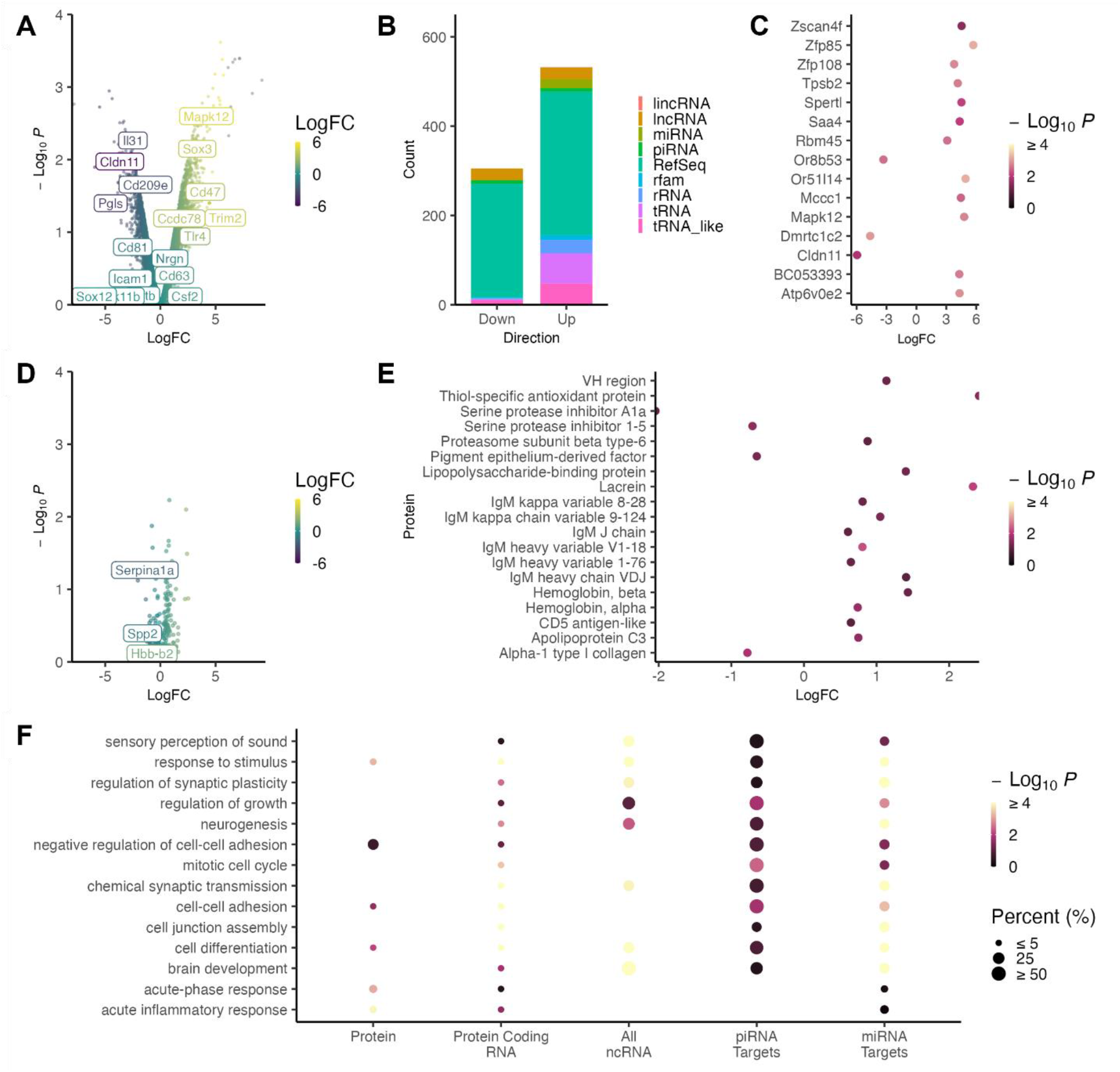
FUS-BBBO alters murine EV RNA and protein load. **A)** Volcano plot of differentially expressed RNA between 1 hour post-treatment and baseline EVs with key protein-coding genes highlighted. **B)** Bar chart of the significantly (p<0.05) up and down-regulated RNA split by type. **C)** Plot of most significantly up and downregulated protein-coding RNA with LogFC of protein expression on the horizontal axis. **D)** Volcano plot of differentially expressed proteins (from mass spectrometry proteomics) between 1 hour post-treatment and baseline EVs with key proteins highlighted. **E)** Plot of most significantly up- and downregulated proteins with LogFC of protein expression on the horizontal axis. **F)** Functional annotation of the 1) significantly upregulated proteins, 2) significantly upregulated protein-coding genes, 3) significantly upregulated non-coding RNA (ncRNA), 4) genetic targets of the significantly upregulated piRNA, and 5) genetic targets of the significantly upregulated micro-RNA (miRNA). Adjusted Kolmogorov-Smirnov p-value magnitude is displayed in color. The size of each dot corresponds to the percentage of annotated genes from that term that are significantly upregulated.

Next, functional annotation was used to identify the biological processes associated with the differentially expressed: a) proteins, b) protein-encoding genes, c) ncRNA, d) piRNA target genes, and e) miRNA target genes. Protein functional annotation maps to immediate and acute inflammatory response, while the protein-coding and non-coding RNA correspond to more long-term responses such as synapse regulation and neurogenesis (Fig. **2F**). Many of the functions associated with the RNA changes are coincident with reported FUS-BBBO increases in neurogenesis [5], proliferation [5], and synaptic remodeling [13]. This leads to the hypothesis of EV response involvement in the neuroimmunotherapeutic responses to FUS-BBBO.

### GW4869 eliminates murine EV concentration increase and reduces inflammatory response

To further elucidate the role of EVs in modulating FUS-BBBO neuroimmunotherapy, we utilize GW4869, a neutral sphingomyelinase inhibitor that is the most widely used agent for blocking EV generation [25] (Fig. **3A**). We studied four groups for this experiment: naïve, naïve+GW4869, FUS-BBBO, and FUS-BBBO+GW4869.

**Figure 3.**
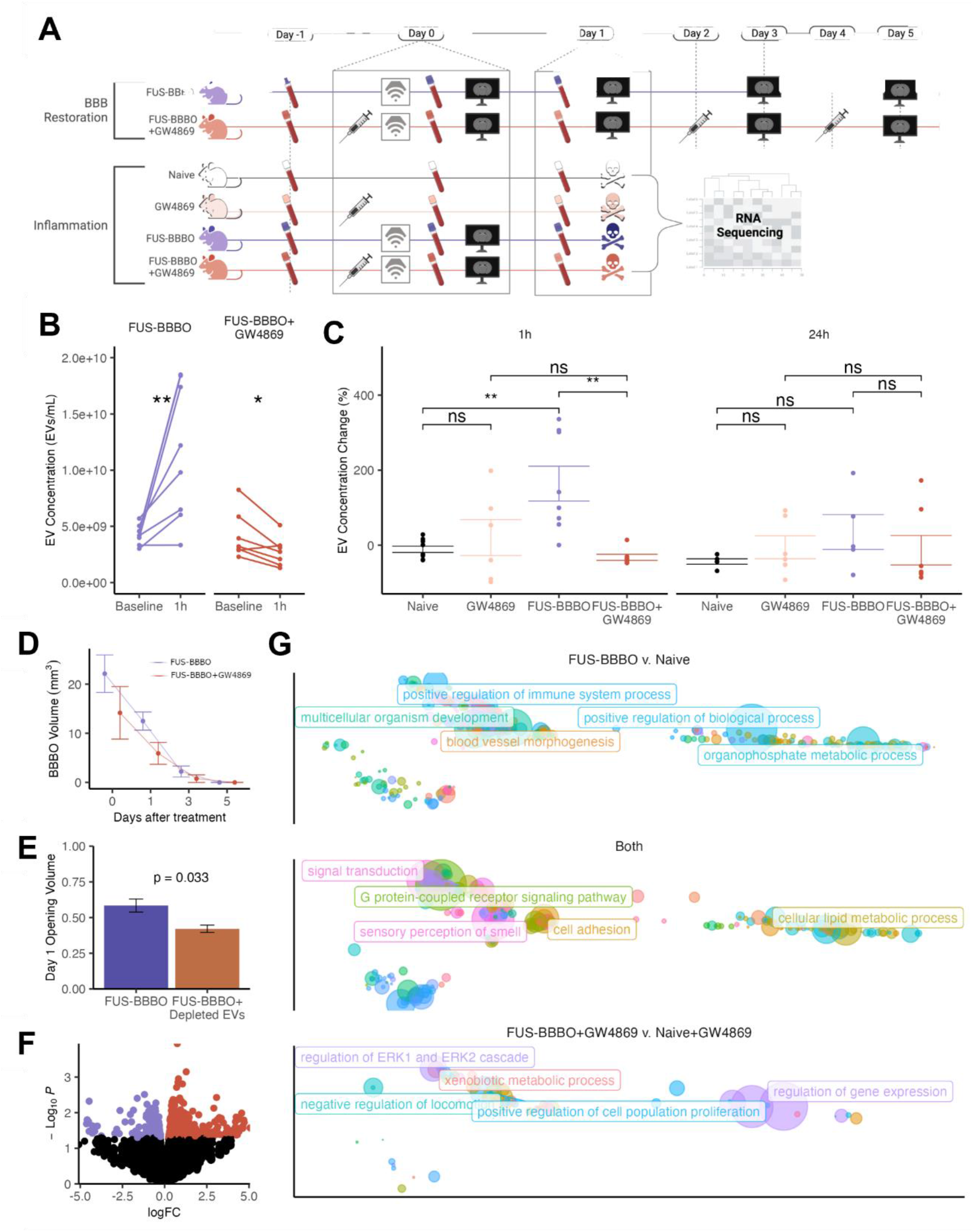
GW4869 eliminates murine EV concentration increase and reduces inflammatory response. **A)** GW4869 study timeline. In the BBB restoration study, sequential MRIs were taken after FUS-BBBO to monitor BBB restoration after treatment with and without GW4869. In the inflammation study, animals were sacrificed 1 day after treatment and the inflammatory response and bulk transcriptome was compared between FUS-BBBO, FUS-BBBO+GW4869, naïve and naïve+GW4869 animals. **B)** EV concentration at Baseline and 1 hour after treatment with either FUS-BBBO or FUS-BBBO+GW4869. Paired t-test was performed for each group. **C)** Percent change of EV concentration 1 hour and 1 day after treatment for all groups. ANOVA followed by Bonferroni post hoc t-tests were performed for each time point. **D)** BBBO volume for the day of treatment and 1 day, 3 days, and 5 days following treatment for both FUS-BBBO and FUS-BBBO+GW4869. **E)** Proportion of original BBBO volume that is still open 1 day after treatment for animals in the restoration study. An unpaired t-test was performed between the two groups. **F)** Volcano plot comparing the profile of FUS-BBBO+GW4869 and FUS-BBBO hippocampi. Significantly expressed (p<0.05) genes are colored. **G)** Significantly upregulated terms from differential gene expression analysis between FUS-BBBO+GW4869 and naïve+GW4869 and FUS-BBBO compared with naïve. Functional ontology terms are clustered by similarity, and the point size shows their significance. Terms appearing in both functional annotations are shown in the “Both” panel with the average significance between the two comparisons.

First, we confirmed that GW4869 successfully eliminated the FUS-BBBO-induced increase in EV concentration. We found that 1 hour after treatment, the animals treated with FUS-BBBO+GW4869 had a statistically lower EV concentration compared to baseline, starkly contrasting with our FUS-BBBO group, which increased in EV concentration by over 100% (Fig. **3B**). Comparing the EV concentration change between our four groups, we see that FUS-BBBO+GW4869 is indistinct from naïve and naïve+GW4869 both 1 hour and 1 day after treatment (Fig. **3C**).

Next, we monitored BBB restoration after FUS-BBBO with and without GW4869. The BBB opening volume of each animal was quantified on Days 0, 1, 3, and 5. On every day of measurement, the animals in the FUS-BBBO+GW4869 had smaller openings than those in the FUS-BBBO group (Fig. **3D**). This difference is particularly significant on day 1 when the BBBO volumes of the FUS-BBBO and FUS-BBBO+GW4869 groups averaged 58% and 42% of the day 0 BBBO volume respectively (Fig. **3E**).

Due to previous literature identifying 1 day as the peak of FUS-BBBO induced inflammation [5], [31], [32], and given the more restored BBB in our FUS-BBBO+GW4869 group at this time point, we performed bulk RNA sequencing on tissue extracted from 1 day after FUS-BBBO for our four treatment groups. Differential gene expression between FUS-BBBO and FUS-BBBO+GW4869 revealed that GW4869 reduced the presence of inflammatory markers, including *IL6* and *CCL4* (Fig. **3F**). In order to account for any effects of GW4869 treatment alone, we performed functional annotation on the differentially expressed genes between FUS-BBBO compared with naïve and FUS-BBBO+GW4869 compared with naïve+GW4869. This revealed increased processes altered in FUS-BBBO without injecting GW4869, including migratory, development, and inflammatory terms. The functions shared by both comparisons are primarily involved with vasculature development (Fig. **3G**).

Overall, we see that GW4869 eliminates the FUS-BBBO-induced increase in EV concentration, decreases the volume of BBB opening, and reduces the number of differentially affected processes after FUS-BBBO, indicating a vital role of EVs within the neuroimmunotherapeutic response to FUS-BBBO.

### Patient extracellular vesicle concentration peaks 1 hour after FUS-BBBO

Six Alzheimer’s Disease Patients underwent FUS-BBBO as part of our group’s phase I clinical trial (NCT04118764) as reported in our group’s clinical paper [29]. All patients had blood drawn immediately prior to treatment (Baseline) and 3 days after treatment. Additionally, the last four patients — P3, P4, P5, and P6 — had blood drawn 1 hour after treatment (Fig. **4A**). All patients apart from P3 had confirmed blood-brain barrier opening; P3 did not have successful opening and thus is considered a FUS-sham subject.

EVs were isolated from each time point for each patient. Western blotting of the isolated EVs from representative Baseline and FUS-BBBO samples confirmed successful EV isolation (Fig. **4B**). Comparing EV concentration from Baseline to 1 hour post-treatment reveals a significant increase in EV concentration (Fig. **4C**) with a near-return to Baseline by 3 days after treatment (Fig. **4D**). Furthermore, the percent increase in EVs 3 days after treatment is correlated with the volume of the blood-brain barrier opening (Fig. **4E**).

**Figure 4.**
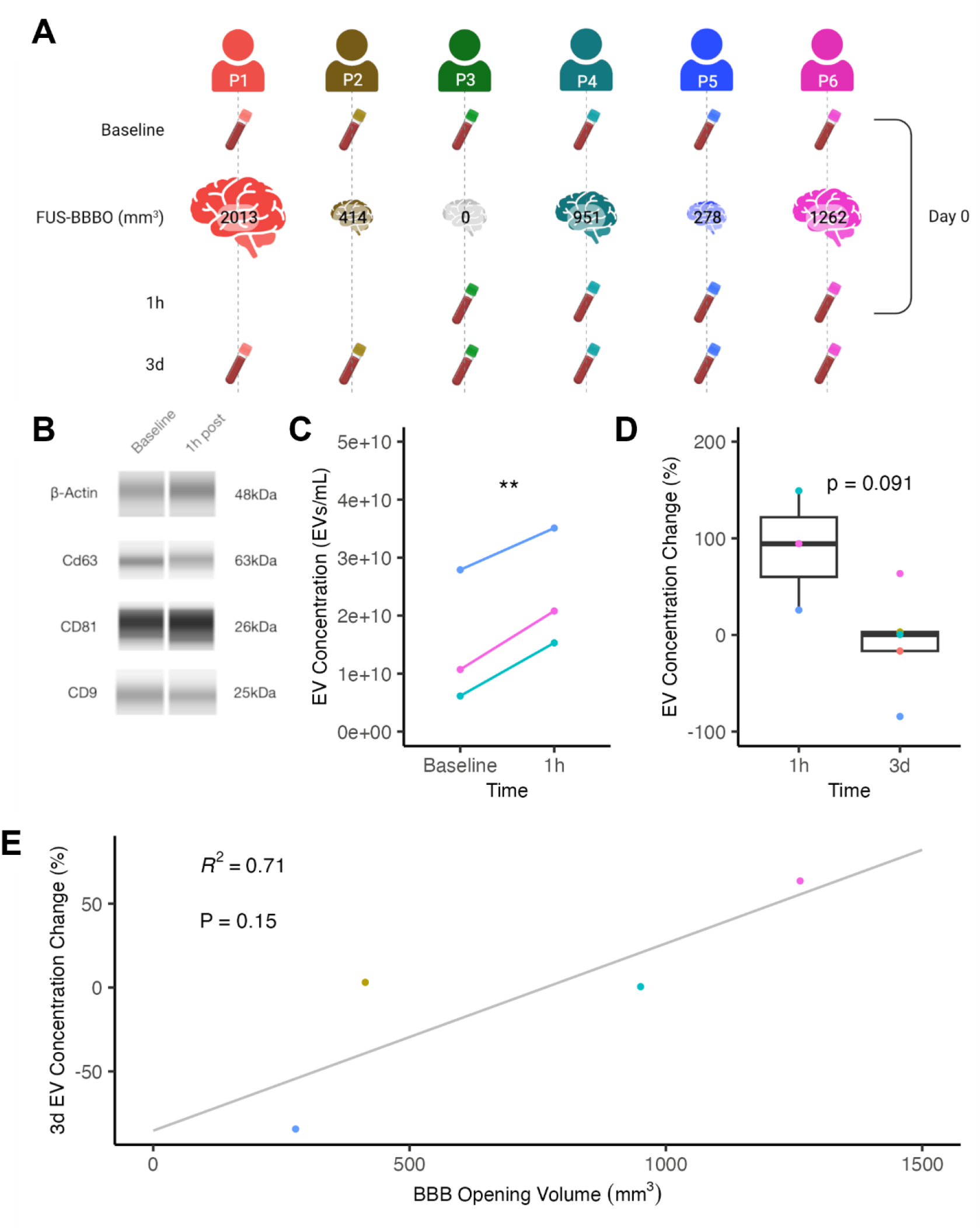
Extracellular vesicle concentration increases after FUS-BBBO in Alzheimer’s Patients. **A)** Schematic of clinical trial with neuronavigation-guided FUS-BBBO, including each patient’s blood draw and opening volume. **B)** Western blots of β-actin and tetraspanins CD63, CD81, and CD9 from isolated EVs at baseline and 1 hour after FUS-BBBO confirm successful EV isolation. **C)** EV concentration at Baseline and 1 hour after treatment for the patients who had successful treatment sessions. **D)** Percent change in EV concentration 1 hour and 3 days after treatment compared to baseline. An unpaired two-tail t-test was performed between the two groups. **E)** Correlation of BBBO volume and the percent change in EV concentration 3 days after treatment. Simple linear regression was performed, and the resulting R2 and p-value are on the chart.

### EV isolation improves FUS-BBBO amplification of neurological biomarkers

To elucidate the utility of EVs in improving liquid biopsy specificity, we quantified several potential AD biomarkers, EV proteins, and other CNS proteins within our patient-isolated EVs, isolated EVs normalized by EV concentration (normalized EVs), and total serum. We find that the EV, normalized EV, and total serum protein concentrations remain mostly unchanged 1 hour after treatment and even decreased compared to baseline. Three days after treatment, the concentration of a number of proteins is significantly increased for both EVs and normalized EVs compared to the total serum, which has no changes in marker concentration for any of the markers (Fig. **5A**). Furthermore, the differences between 1 hour and 3 days log fold change (LogFC) in protein content are more statistically distinct for the EVs and normalized EVs than the total serum (Fig. **5B**).

Finally, in our clinical study, we compared the MRI-based blood-brain barrier opening volume to the LogFC of select biomarkers 3 days after treatment [29]. Further analysis of this data indicates that the level of the AD biomarkers is positively correlated with BBBO in serum and EV content but not normalized EV content (Fig. **5B**). This indicates that the increase in biomarker detection is due to more EVs being released, and not because each EV has higher biomarker concentration. Overall, our results from the AD clinical trial reveal the potential of FUS-BBBO in enhancing EV-based biomarker detection sensitivity and specificity for liquid biopsy.

**Figure 5.**
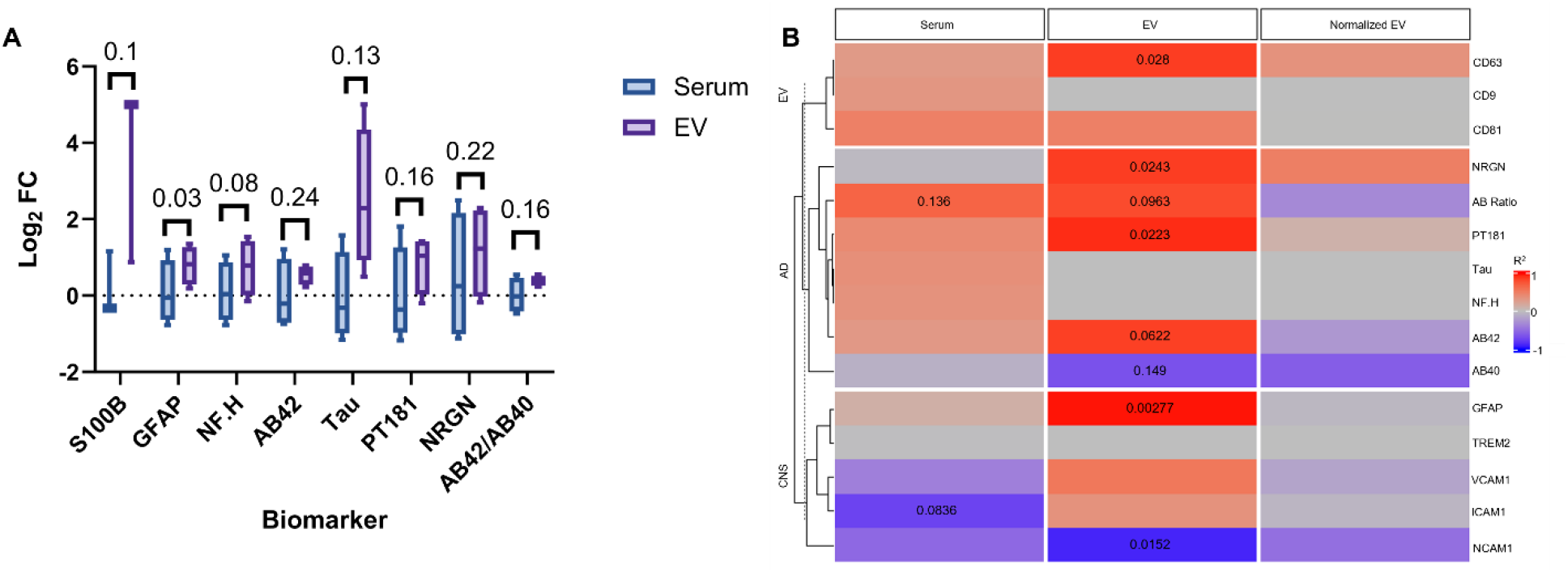
EV isolation further improves FUS-BBBO amplification of neurological biomarkers. **A)** Comparison of the FUS-BBBO induced increase in neurological biomarker detection in total serum (blue) and isolated EVs (purple) with a paired t-test performed for each biomarker. **B)** Correlation between FUS-induced volume of opening and the LogFC change of biomarkers in total serum, isolated EVs and isolated EVs normalized to the EV concentration (Normalized EV). Simple linear regression was performed for each condition, and R2 and p values are displayed on the heatmap.

## Discussion And Conclusion

Focused ultrasound blood-brain barrier opening (FUS-BBBO) has been studied as a neuroimmunotherapeutic and a method of improving liquid biopsy specificity for neurological disorders [4], [6], [11]. Due to the dual role of extracellular vesicles (EVs) in modulating the neuroimmune system [18], [19] and as an emerging biomarker [20], [33], we aimed to identify the effect of FUS-BBBO on EVs isolated from mouse and patient serum.

We first identified a significant increase in mouse EV concentration 1 hour after treatment coincident with RNA changes associated with the EV-dependent neuroimmunotherapeutic effects of FUS-BBBO such as neurogenesis, synaptic pruning, and barrier maintenance [5], [6], [16], [34], [35]. Eliminating the EV concentration increase with GW4869 resulted in reduced blood-brain barrier opening volume and inflammation, indicating the contribution of the EVs to FUS-BBBO inflammation and opening volume. The immune response to FUS-BBBO has spearheaded debate about the method’s safety, so the ability to control and mitigate FUS-BBBO-induced inflammation provides an exciting new avenue, mainly when FUS-BBBO is used purely as a drug delivery tool [32]. Future work will include investigating the neuroimmunotherapeutic capacity of FUS-BBBO with depleted EVs because those benefits may require a complete neuroimmune response.

Secondly, we identified a significant increase in Alzheimer’s Disease (AD) patient EV concentration 1 hour after treatment that remained dependent on BBBO volume 3 days after treatment. This was accompanied by increased AD biomarker detection specificity in isolated EVs compared to total serum, which was also correlated with the volume of BBBO as shown in our clinical study [29]. These results highlight the potential of EVs to be used as accurate and specific markers for neurological diseases, and the ability of FUS-BBBO to amplify their detection in a non-invasive way.

However, in order to be implemented for improving biomarker specificity, there needs to be a method of differentiating between the changes in biomarker concentration due to the treatment itself and those due to the disease. Other groups have addressed this challenge by utilizing binary biomarkers such as cfDNA, which are present in the case of disease or otherwise completely absent [7], [9]. This becomes much more complex with diseases such as AD, where many of the proposed biomarkers are CNS proteins that are always in the CNS albeit in different concentrations [20], [33]. Future research must include a normalization between BBBO and biomarker concentration that can be used to identify a BBBO-induced concentration change compared to a disease-induced concentration change.

Overall, this study presents the first preclinical and clinical evidence of FUS-BBBO increasing EV concentration and altering EV content, which has implications in both the mechanism of FUS-BBBO neuroimmunotherapy and the optimization of FUS-BBBO-induced amplification of neurological disease biomarkers.

## Abbreviations

AD: Alzheimer’s Disease
BBB: Blood-brain barrier
cfDNA: Cell-free DNA
CNS: Central nervous system
EV: Extracellular vesicle
FUS: Focused ultrasound
FUS-BBBO: Focused ultrasound-mediated blood-brain barrier opening
GFAP: Glial fibrillary acidic protein
HRP: Horseradish peroxidase
LC/MS ESI-TOF: Liquid chromatography/mass spectrometry electrospray ionization time-of-flight
LogFC: Log fold change
MB: Microbubble
miRNA: Micro-RNA
MRI: Magnetic resonance imaging
ncRNA: Non-coding RNA
NTA: Nanoparticle tracking analysis
PBS: Phosphate-buffered saline
piRNA: Piwi-interacting RNA.

## Acknowledgements

This study was supported in part by the National Institutes of Health under Grants R01AG038961, R01EB009041, and R56AG038961, and the Focused Ultrasound Foundation. FNT was also supported by the Onassis Foundation under contract number F ZT 072-1/2023-2024. Some figures were created with BioRender.com. The authors wish to thank UEIL members Rebecca Noel, Ph.D., Daniella Jimenez, B.S., Samantha Gorman, B.S., and Nancy Kwon M.S. for their support and insightful scientific discussions.

## Contributions

ARKS, AJB, and EEK designed the study and the methodology. ARKS, AJB, and MRD conducted in *vivo* mouse studies. KL and SB collected and processed blood samples from the clinical study. ARKS, FNT, and MRD processed EV results and made figures. ARKS, FNT, and EEK drafted and revised the manuscript. EEK acquired funding and provided resources for the study. All authors contributed to discussions and reviews of the manuscript.

